# Of Problems and Opportunities – How to Treat and How to Not Treat Crystallographic Fragment-Screening Data

**DOI:** 10.1101/2022.06.01.492756

**Authors:** Manfred S. Weiss, Jan Wollenhaupt, Galen J. Correy, James S. Fraser, Andreas Heine, Gerhard Klebe, Tobias Krojer, Marjolein Thunnissen, Nicholas M. Pearce

**Affiliations:** Macromolecular Crystallography, Helmholtz-Zentrum Berlin, D-12489 Berlin, Germany; Department of Bioengineering and Therapeutic Sciences, University of California San Francisco, San Francisco, CA 94158, USA; Institute of Pharmaceutical Chemistry, Philipps University Marburg, D-35032 Marburg, Germany; MAX IV Laboratory, Lund University, PO Box 118, S-221 00 Lund, Sweden; Department of Chemistry and Pharmaceutical Sciences, VU Amsterdam, 1081 HZ Amsterdam, The Netherlands

**Keywords:** group depositions, fragment-screening, low-occupancy ligands, PanDDA, compositional heterogeneity, conformational heterogeneity

## Abstract

In their recent commentary in Protein Science, Jaskolski *et al*. analyze three randomly picked diffraction data sets from fragment-screening group depositions from the PDB and, based on that, claim that such data are principally problematic. We demonstrate here that if such data are treated properly, none of the proclaimed criticisms persist.

## INTRODUCTION

In their recent commentary in Protein Science^1^, M. Jaskolski and colleagues suggest that group depositions arising from large crystallographic screening campaigns to the Protein Data Bank wwPDB^2^ are problematic and pose a serious threat to the integrity of the database. They mention that the group deposition models “*do not conform to the quality standards expected*”, that they “*confuse most biomedical researchers*” and that they “*degrade the PDB integrity*”. In order to overcome this, they postulate that such group depositions should either be “*clearly marked*” or they “*should be relocated from the PDB into a separate database*”. Admittedly, some of the concerns of Jaskolski and colleagues are justified. However, PDB procedures and mechanisms, including optimized guidelines for group depositions are also rapidly evolving. Still, as with all new techniques, they will need some time to mature. In particular, standards for group depositions from fragment-screening campaigns are yet far from clear cut. Consequently, group depositions are often not adequately marked within the PDB and can even contain different data items. These attributes can make it difficult for the some PDB users, especially for those without an extensive structural biology background, to analyze and interpret these structures properly.

Here, we would like to caution that the arguments advanced by Jaskolski *et al*. are perhaps too one-sided. They contrast refined data from single structure determinations with group depositions where the relevant information is spread over many individual data sets and they conclude that the former are *per se* “*better*” and meet expected quality standards, particularly of placed ligands more precisely, whereas the latter are likely to suffer from reduced quality. We would like to assert, that these conclusions do not accurately reflect the complexity of these data and may, in fact, in our opinion hinder progress in the field of structural biology, in particular of drug discovery programs, in which these new methods are already widely applied. Commentaries that underestimate the knowledge of PDB users, that ignore the opportunities present in heterogeneous crystallographic data, and that miss out on chances for education on structure quality do more harm than good to structural biology. In the end, it may even sow distrust among end-users with respect to new methods to analyze valid primary data. Contrasting their rather conservative view of “a single dataset, a single structure, a single interpretation”, we would like to inspire now a more positive and optimistic look at recent developments in particular in crystallographic screening campaigns. We envision a near future where many datasets can be collectively analyzed, bringing about an ensemble view of macromolecular structure that fully embraces both conformational and compositional heterogeneity in the underlying data.

Undoubtedly, the last decade has seen tremendous changes in macromolecular crystallography at synchrotron beamlines. Improvements to sample handling automation, detector speed and sensitivity, and on-the-fly data processing have made it possible to record and analyze large numbers of diffraction data sets^3,4^. These advances have been foundational to the use of “crystal soaking with small fragments followed by crystallography” as an extremely sensitive “binding assay” in fragment screens^4–8^. With further medicinal chemistry, the identified binders in such fragment screens may then be developed by rational design concepts into stronger binding compounds and ultimately into drugs^9^. While it was plainly impossible to record several hundreds of diffraction data sets within a manageable amount of time ten years ago, it is now almost routine within just 24 hours of beam time^4^ at several synchrotron sites around the world^4–7^. Industrial pharmaceutical research has heralded such experiments much earlier than the academic sector, however, due to intellectual property concerns, few of those results have been made publicly available. As academic efforts have intensified, fragment screens are reported more frequently in the recent literature^10-12^, and their results are deposited as a batch in the wwPDB^2^. Without any doubt, these data will have tremendous impact on health sciences and at the same time help pave the ground for new and essential techniques in data handling and evaluation.

Crucially, these developments have been enabled by diverting from the traditional paradigm of “a single dataset, a single structure, a single interpretation”. A typical crystallographic fragment screening campaign comprises hundreds of individual diffraction data sets. In order to efficiently analyze the entire screen as one overall data set, Pearce and van Delft developed the Pan-Dataset Density Analysis (PanDDA) procedure^13^. PanDDA exploits the fact that the hundreds of structures are all nearly identical and exhibit correlated signals and errors. PanDDA works by aligning individual density maps and identifying spatially contiguous voxels with signals outside the background distribution. The background distribution can then be subtracted, resulting in an “event map”, which is the most crucial read-out of a fragment screening experiment. The event map represents the primary evidence for the presence of a ligand, allowing the identification and modelling of low-occupancy ligands that are often elusive to classical (mFo-DFc)-difference-density based approaches. By inherently embracing the idea that the contents of the crystal are *compositionally* heterogeneous, in this case possibly as a result of the applied soaking procedure, this procedure is much more sensitive than treating the individual datasets in isolation and analyzing each for differences relative to a single “apo” dataset. As a consequence, individual fragments are often identified, despite having such low occupancy in the crystal that they are imperceptible in both the original electron density maps and by traditional reciprocal space difference map approaches.

## RESULTS

Jaskolski *et al*.^1^ are right to note that such a new approach with reporting and analyzing batch-data does not conform to some of the classical PDB expectations and that they cannot and should not be treated in the same way. However, what they do is that they pull out three individual coordinate sets from three different group deposition entries and look at them using the classical difference electron density map approach. It is therefore not surprising that they encounter the problems they reported. In the first case (PDB-Id 5RTL^12^) they do not observe a good agreement of low-occupancy ligand density with the associated (mFo-DFc) difference electron density map (**Figure 1A**) and question the presence of the ligand at all. However, if Figure 1a of Jaskolski *et al*. is contrasted by the PanDDA generated event map of PDB-Id 5RTL, the evidence for the presence of the ligand is clearly there. In their second example (PDB-Id 5RDH^10^), Jaskolski *et al*. report that the ultra-high resolution of the data set does not match the expected quality indicators of the structure (**Figure 1B**). They note that the ultra-high resolution may confuse non-expert users of the model and potentially lure them into using it as a reference model. However, a closer look at the data processing statistics reveals that the ultra-high resolution is just a consequence of a hitherto undiscovered problem, leading to a faulty resolution cutoff during the automated data processing step^14^. Simple reprocessing yielded a data set to 0.93 Å resolution and a free R-factor after auto-refinement of 13%, which is perfectly in the expected range (**Figure 1C**). The third case that is reported (PDB-Id 5RFB^15^) is similarly as in the first case where the ligand is in the focus (**Figure 1C**). Also here, the presence of the ligand is clearly supported by the respective event map. In this context, we fervidly refute the statement of Jaskolski *et al*. that “*No useful conclusions can be derived by PDB users from this ligand* …”. Again, if the data are looked at properly^16^, accounting for the compositional heterogeneity and the fact that the ligands are low occupancy, none of the described criticisms persists (**Figure 1**).

**Figure 1.**
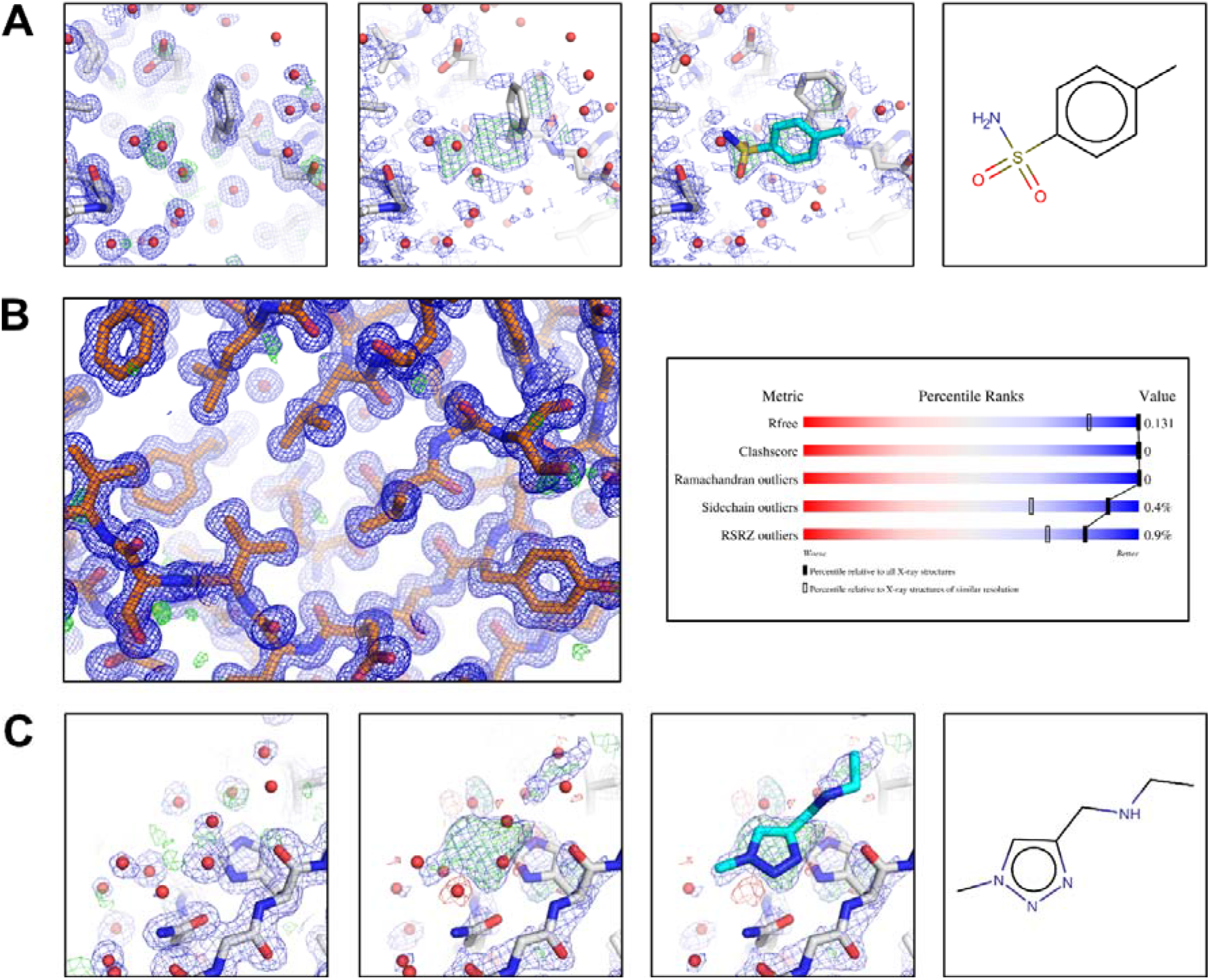
**A)** Ligand identification for PDB-Id 5RTL. Left panel: auto-refinement model, (2mFo-DFc) map, contoured at 1.0 σ (blue), (mFo-DFc)-difference map contoured at 3.0 σ (green/red, similar to Figure 1a of Jaskolski *et al*.,) Middle panels: PanDDA Z-map (contoured at Z=3, green/red) and PanDDA event map contoured at 1.0 σ (blue), with auto-refined model and ligand placed (=bound state) model, respectively. PanDDA event map coefficients are available from the PDB in the deposition’s structure factor CIF. Right panel: chemical structure of the ligand, ZINC388056. **B)** PDB-Id 5RDH after reprocessing. Left panel: autorefined model and (2mFo-DFc) map, contoured at 1.0 σ (blue), (mFo-DFc)-difference map contoured at 3.0 σ (green/red). **C)** Ligand identification for PDB-Id 5RFB. Left panel: auto-refinement model, (2mFo-DFc) map, contoured at 1.0 σ (blue), (mFo-DFc)-difference map contoured at 3.0 σ (green/red, similar to Figure 1a of Jaskolski *et al*., 2022) Middle panels: PanDDA Z-map (contoured at Z=3, green/red) and PanDDA event map contoured at 1.0 σ (blue), with auto-refined model and ligand placed (=bound state) model, respectively. Right panel: chemical structure of the ligand, Z1271660837.

## DISCUSSION

Unfortunately, based on these three cases, Jaskolski *et al*. come to the general conclusion that models from such group depositions will contaminate the PDB and that they should be deposited differently in a distinct resource, potentially even outside the PDB. Obviously, we agree on the notion that group depositions ought to be represented differently from single crystal data sets. Instead of demanding such “offending” models to be removed from the data bases such as the PDB, it seems that the real question to be asked is: “*how* should low-occupancy ligand structures and multi-state crystal structures be presented in the PDB?” At the moment, there seems to be no satisfactory way to deposit full multi-state ensembles. With proper data management and curation tools and data sets accompanied by metadata for handling multi-state models in place in the PDB, however, this issue could be immediately solved allowing depositing of full sets of crystallographic data and also clearly presenting the “states of interest” to the (non-specialist) user. Simply banning structures with low-occupancy ligands would almost certainly negatively interfere with future developments in structural biology.

The idea that depositing data beyond what might be of biological interest today is reflected in the instructive example of the surprising usefulness of Structural Genomics. About 25 years ago, Structural Genomics engaged in the massive determination of macromolecular structures with no particular biological question associated with the vast majority of the targets. Back then, some people in the field argued: “*These semi-automatically determined structural genomics structures are worse than the handmade structures*”, that “*Structural Genomics is like stamp collecting*^*17*^ *with no scientific meaning*” and so forth. Irrespective of that, Structural Genomics pushed forward and thousands of these structures ended up in the PDB, many of them without an associated publication. What is more, Structural Genomics triggered major advances not only in algorithms and automation, but also solutions to less obvious problems such as data standardization, paving the way for next-generation developments like crystallographic fragment screening. But the benefits appear even far more wide-ranging: two decades later, clever scientists from way outside structural biology, leveraged these diverse coordinate sets to solve the long sought after sequence-to-structure prediction problem. It is quite likely that, without Structural Genomics, there would be no AlphaFold2^18^.

So here we are again… “*Thou shalt not contaminate the PDB*”, we can hear the gatekeepers of the holy structure roar, “*with your fragment-screening data sets*”. But then we may simply counter this with “why not?”, as there will always be scientists who can make use of our data for novel developments and methods in a much cleverer way than we can currently imagine. Likely people will soon invent smart ways to efficiently extract all aspects of *conformational* as well as of *compositional* heterogeneity out of all these data sets. In doing so they might even “solve” protein conformational dynamics or protein-ligand binding prediction, much the way AlphaFold2 has “solved” protein structure prediction. As long as the data is there, let’s embrace it and make it available!

## METHODS

### PDB-Id 5RTL

Diffraction data and coordinates for PDB-Id 5RTL^12^ were retrieved from the PDB. Auto-refinement from the intermediate DIMPLE^19^ step yielded a model and a set of model phases that approximately reproduce the findings by Jaskolski *et al*.^1^ (see Figure 1a of Jaskolski *et al*. and **Figure 1A**, leftmost panel). The PanDDA event map (available from the PDB under PDB-Id 5RTL) and the PanDDA Z-map (provided by the authors) clearly indicates the presence of the ligand ZINC388056 (**Figure 1A**, third panel from the left).

### PDB-Id 5RDH

The diffraction images corresponding to PDB-Id 5RDH were retrieved from storage and reprocessed using XDSAPP^14^ using all default parameters. This resulted in a data set extending to 0.93 Å resolution. Automatic refinement using fspipeline^7^ yielded a coordinate set, which conforms very well to all conventionally applied refinement and model statistics (**Figure 2B**).

### PDB-Id 5RFB

Diffraction data and coordinates for PDB-Id 5RFB^15^ were retrieved from the PDB. Auto-refinement from the intermediate DIMPLE^19^ step (D Fearon and F von Delft, personal communication) yielded a model and a set of model phases that approximately reproduce the findings by Jaskolski *et al*.^1^ (see Figure 1c of Jaskolski *et al*. and **Figure 3A**, leftmost panel). In contrast, the PanDDA event map (available from the PDB under PDB-Id 5RFB) and the PanDDA Z-map (D Fearon and F van Delft, personal communication) clearly indicates the presence of the ligand (**Figure 3C**, second and third panel from the left, respectively).

## ACKNOWLEDGEMENTS

We would like to thank Daren Fearon and Frank von Delft (Diamond Light Source, UK) for providing the map and model files used to generate **Figure 1C**. We are also grateful for Hans Wienk (NKI Amsterdam, The Netherlands) for critically reading the manuscript and for many textual improvements.

